# Machine-learning algorithm for identifying and predicting amyotrophic lateral sclerosis causal mutations

**DOI:** 10.1101/2022.03.27.485996

**Authors:** Serveh Kamrava, Ali Sahimi, Justin Ichida, Muhammad Sahimi

**Affiliations:** Department of Chemical Engineering and Materials Science, University of Southern California, Los Angeles, California 90089, USA; Department of Stem Cell Biology and Regenerative Medicine, University of Southern California, Los Angeles, California, 90089 USA

## Abstract

We propose a machine learning (ML) method to classify ALS–causative and non–ALS–causative variants based on 24 variables in five different datasets. The proposed ML method classifies the five datasets with very high accuracy. In particular, it predicts the ALS variants with 100 percent accuracy, while its accuracy for the non-ALS variants is up to 99.31 percent. The trained classifier also identifies the nine most influencial mutation assessors that help distinguishing the two classes from each other. They are FATHMM_score, PROVEAN_score, Vest3_score, CADD_phred, DANN_score, meta-SVM_score, phyloP7way_vertebrate, metaLR, and REVEL. Thus, they may be used in future studies in order to reduce the time and cost of collecting data and carrying out experimental tests, as well as in studies with more focus on the recognized assessors.

## I. INTRODUCTION

Amyotrophic lateral sclerosis (ALS) is characterized by a progressive loss of motor neurons and denervation of muscle fibers, which result in muscle weakness and paralysis. With an incidence of 2.7 cases per 100,000 individuals [1], it is a common disorder that is more prevalent in the later years of life. There is currently no medical cure for the disease, and only two drugs, riluzole [2] and radicava [3], slow progression of the disease moderately. ALS pathology manifests a specific degeneration of the upper and lower motor neurons, leaving many other cell types unimpaired. The disease is multifactorial, as multiple processes play a role, including protein aggregation, excitotoxicity, and RNA processing impairments [4]. A broad range of therapeutic strategies have been attempted in rodent models of ALS, but positive results have not been frequently reproduced in human trials [5].

Approximately 90 percent of ALS cases are sporadic (sALS), with the remaining 10 percent of the cases having a family member with the disease (fALS) and, thus, they likely inherited a disease-causing mutation. ALS is mostly viewed as a monogenic disease, with SOD1 mutations accounting for approximately 10-20 percent of all familial cases. More than 100 different mutations in the SOD1 gene cause ALS with varying severity and penetrance [6]. The genetic variation that explains the largest number of ALS cases is the GGGGCC repeat expansions in gene chromosome 9 open reading frame 72 (C9ORF72), also known as the C9ORF72 hexanucleotide repeat expansion, as the repeat expansion causes ALS in about 23.5 percent of fALS [7]. The intronic hexanucleotide repeat expansion in C9ORF72 is also the most common cause of ALS and frontotemporal dementia (FTD), as it accounts for over 50 percent of ALS in northern Europe and 10 percent of cases worldwide.

About 70 percent of fALS and 15 percent sALS can be explained by known ALS mutations [7]. It is difficult to identify new ALS–causing mutations because each etiology seems to be rare, and ALS is already a relatively rare disease with only 30,000 patients in the United States. While it is possible to test candidate causal mutations by CRISPR-Cas9 gene editing in cell or animal models, the average number of rare variants per individual far exceeds the number of variants that one can test through CRISPR editing. In addition, the heredity of sALS due to common variants is predicted to be about 18 percent [8], implying a strong basis for further genetic investigation into ALS, both in fALS and in sALS. Interestingly, approximately 30 genes have been associated with ALS, and the mutations can cause the disease by gain-of-function (GOF), as well as by loss–of–function (LOF), mechanisms and contribute to multiple pathways. Moreover, as fALS and sALS are clinically indistinguishable, common disease mechanisms may be expected.

The clinical presentations are, however, quite variable between patients, including large differences between age of the onset - from early 20s to 70s - site of the onset - limbs vs. bulbar - and progression rate with survival of less than 6 months to 10 percent of the patients surviving longer than 10 years. Some mutations are correlated with particular ALS phenotypes, e.g., mutations in SETX frequently cause young onset ALS [9-14]. As ALS is a multifactorial disease where many biological mechanisms contribute to motor neuron death, and it is not yet clear which mechanisms are crucial to effectively treat (specific) ALS patients, it is important to be able to study the pathophysiology induced by each new ALS mutation.

Two noteworthy papers should be mentioned here. As a new screening strategy, Kenna *et al*. [15] carried out entire-exome analyses of 1,022 index fALS cases, and 7,315 controls. They carried out gene-burden analyses, trained with established ALS genes and identified a significant association between the LOF NEK1 variants and fALS risk. Hu *et al*. [16] used the current understanding of disease genetics in order to train ML models and to predict novel genetic factors associated with the various diseases, although they did not use what we propose in this paper. They developed DGLinker, a webserver for the prediction of novel candidate genes for human diseases, given a set of known disease genes.

In this paper we put forth a hypothesis, and describe a machine-learning (ML) algorithm for testing it. We hypothesize that ALS-causal gene variants can be predicted and identified by contrasting them with non-disease causal ones, and that identification of the ALS-causal variants is possible and predictable. Clearly, if the hypothesis is true, its implication would be imperative for the development of novel therapeutic strategies, especially if gene-correction or gene-editing strategies become advanced enough for use in patients with motor neuron disease. Our proposed ML algorithm provides classification of the data with extremely high accuracy, which to our knowledge, has not previously been used for ALS or any other disease with a similar genetic architecture.

## II. THE DATA

Variants were curated from the collection held by the Amyotrophic Lateral Sclerosis Online database [17]. Each variant entry included the original article in which the method for its discovery is described, which were utilized to filter the entries and keep those whose described variants were confirmed to have been inherited with full segregation, i.e., those for which both the diagnosis of disease and sequencing of the variant were confirmed in at least two consecutive generations in the proband’s family. Additional papers were obtained from PubMed and other sources. Through examination of the results indicated that all the collected variants were nonsynonymous single nucleotide polymorphism (non-SNPs), located in the exonic regions. Twenty two variants were collected that matched the criteria for high likelihood of causality that we set. We refer to this as dataset number one.

Control variants that were designated as “non-ALS” were curated from supercentenarian patients in the previous sequencing data studies [18]. Variants were filtered out in multiple increments, and the filtered and non-filtered groups were statistically compared to each other to see whether there are significant differences between their mean allele frequencies and quality control scores. Initially, the data were annotated using the ANNOVAR software that provides each variant with pathogenicity scores from multiple algorithmic softwares, such as FATHMM, PolyPhen, MetaSVM, and others. Variants were also annotated using an R script with REVEL, whose data file was downloaded from an open source [19].

The initial dataset of 359,292 variants from supercentenarian individuals was filtered into lower-frequency group (with frequency less than 0.05), which comprised of 18,639 cases, and a higher-frequency one with the remaining 340,550 cases, in order to remove the more highfrequency variants that are common to the population. The lower-frequency group was filtered from the overall set contained variants whose population frequency was higher than what is expected for ALS variants. The primary reason that such variants were still included was having an adequate number of data points for the training and testing tests. Next, the lower frequency variants were removed if they were synonymous, intronic, and insertion-deletions (8,369 variants), leaving only entries with the SNPs (10,271 variants) to match the characteristics of the ALS variants, and to be properly annotated.

The remaining non-ALS variants served as the basis for deriving five differing control datasets. One dataset, referred to as A, contained variants with the top 5000th score and above, including repeated scores (4,985 variants). Another one, dataset B, consisted of the 90th percentile and above (1473 variants), while dataset C included 958 variants that were present within at least two different individuals. Two datasets, D and E, contained variants from multiple individuals that overlapped with the top 5000th and top 90th percentile sets (622 and 237 variants, respectively).

We then added all the entries from dataset 1 to each of the five non-ALS datasets, in order to generate five different testing and training sets. The resulting five datasets - ALS and non-ALS combined - were annotated using ANNOVAR that provides each variant with pathogenicity scores from multiple algorithmic softwares, including FATHMM, PolyPhen, MetaSVM, and others. Variants were also annotated using an R script with REVEL and MCAP v.1.3, whose data files were obtained from their respective home websites. Before utilizing them in the training of the ML algorithm to test its accuracy, all variants with incomplete pathogenicity scores were removed, as they could not be used with the algorithm if they did not contain scores in all the variables.

After training the algorithm, a single “case” patient sample from the Coriell Institute, which was whole-exome sequenced, was introduced. Genetic data from this patient were filtered through the bcftools pipeline to limit the list to unique variants. Variants were then annotated and filtered in the same manner as the above-described methodology to maintain only SNPs with full annotation present. These variants were introduced to the ML and those that were marked as “ALS-causal” were noted.

## III. COMPUTATIONAL APPROACH

We use a highly-accurate classifying technique that, based on the genetic information, distinguishes the ALS cases from healthy patients, and provides deep understanding of the changes in the variables that may cause fALS. We should, however, point out that while there are extensive data for non-ALS cases, the ALS data are very limited. In such cases one has *class imbalance*, a well-known issue in ML. The aim is, of course, to have high accuracy for the minority class, implying that while having overall high classification accuracy is important, it is even more important to achieve the highest precision for classifying the minority class.

Various techniques that have been used to address imbalanced class problems are either based on data sampling or boosting algorithms. The latter approaches improve the classification predictions by training a sequence of weak models, each compensating the weaknesses of its predecessors, and are based on either reweighting of the data, or resampling, or both. Those that are based on reweighting of the data assign higher weights to misclassified samples, which in most cases are from the minority class. Next, the data with modified weights are fed to the base learner and classification is again carried out. This is repeated iteratively during which new weights are assigned, with the ultimate goal being to classify the input data correctly [21-23]. Among the most common boosting methods that have been developed based on the RUS technique is the RUSBoost algorithm [20,23-25], which randomly adjusts the class distribution in the dataset until a proper class is generated. The algorithm is faster and simpler than the boosting methods that are based on over-sampling and, relative to other boosting algorithms, it requires smaller new training dataset [23].

## IV. THE RUSBOOST ALGORITHM

We have only two possible classes of data, 1 ≡ ALScases and 0 ≡ non − ALScases. In RUSBoost algorithm, described below, *x*_*i*_ ∈ {*X*}, with {*X*} being the input, and *y*_*i*_ ∈ {*Y*} with *Y* representing the output class (which are either 1 or 0). The purpose of the algorithm is to evaluate the “class” or estimate the “probability” *h*_*m*_(*x*_*i*_, *y*_*i*_) → [0, 1]. Initially, the weight of each sample is set to be 1*/N*, where *N* is the number of the training cases. One has *M* weak hypotheses, i.e., batches of input and output data with their own associated weights and, therefore, the algorithm is iterated *M* times. During each iteration various weak hypotheses are selected and tested and the errors in the classification prediction associated with the hypotheses are calculated. Then, the weight of each batch is adjusted such that misclassified examples have their weights increased, whereas the correctly-classified examples have their weights decreased. Therefore, subsequent iterations of boosting will generate hypotheses that more likely correctly classify the previously mislabeled examples.

In practice (see below), one first applies the RUSBoost algorithm to remove samples from the majority class until a certain pre-set percentage *D* of the new temporary training dataset 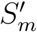 with new weight distribution 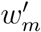 belongs to the minority class. Next, 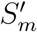 and 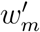 are passed to the next weak learner, the weak hypothesis *h*_*m*_ is generated, and the pseudo-loss *ϵ*_*m*_, as well as the weight update parameter *β*_*m*_, are computed. Then, the weight distribution for the next iteration, *w*_*m*+1_, is updated and normalized. After *M* iterations, a weighted “vote” of the weak hypotheses forms the final hypothesis, *H*(*x*), i.e., the selection of data with the weights that generates the minimum error.

### Algorithm 1 RUSBoost (RUS embedded in AdaBoost)

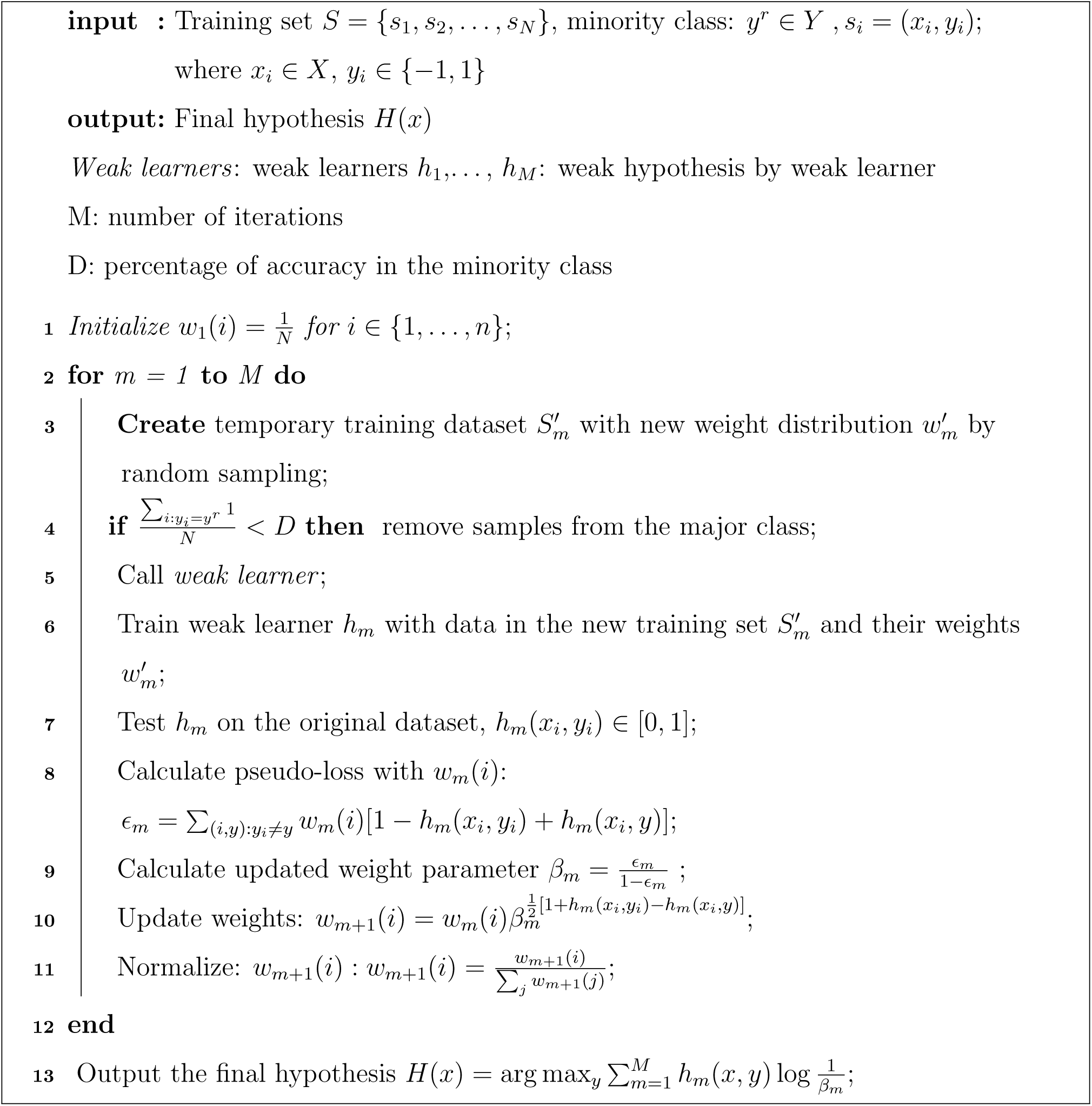

### Performance Metrics

To evaluate the accuracy of the algorithm, we utilize what is called confusion or error matrix, a widely used performance metric for the boosting methods that determines the accuracy of training of the weak learners. The confusion matrix evaluates separately the training accuracy of the two classes, as well as a combination of the two [22]. The classifier is applied to a set of examples for the testing, after which the confusion matrix is constructed. The correctly classified ALS data are placed in the algorithm’s true positive (TP) cell, while the incorrectly classified portion falls into the false negative (FN) cell. Similarly, the correctly classified non-ALS data are labeled true negative (TN), while the incorrectly classified part of this class of data constitutes the false positive (FP) group.

**Table 1:**
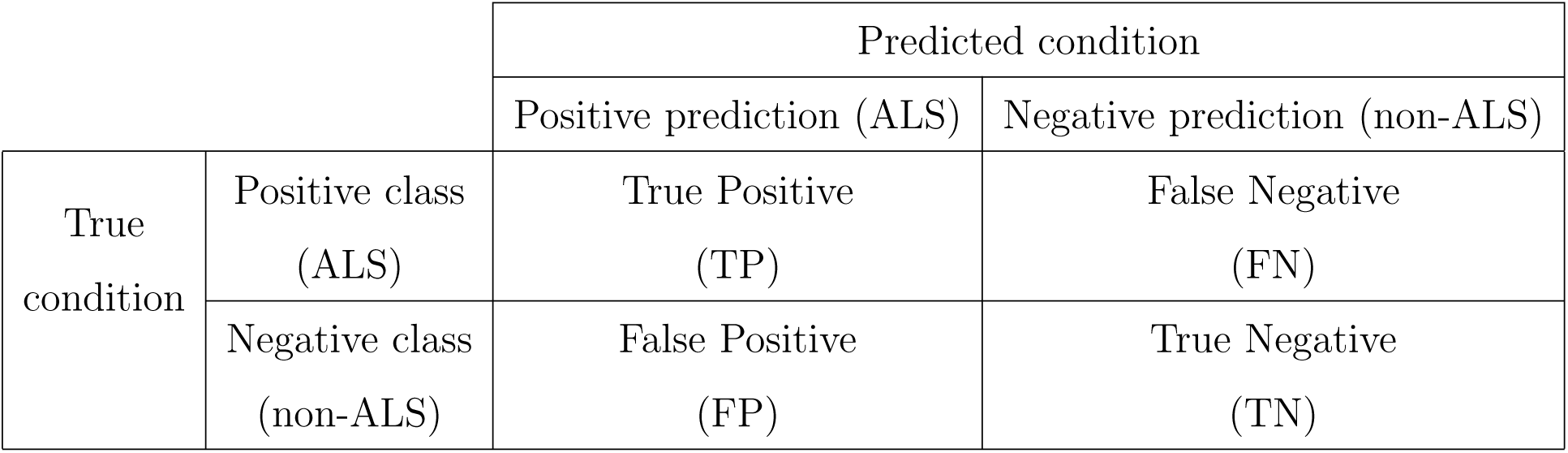
The confusion matrix

|  |  | Predicted condition |  |
| --- | --- | --- | --- |
|  |  | Positive prediction (ALS) | Negative prediction (non-ALS) |
| True condition | Positive class (ALS) | True Positive (TP) | False Negative (FN) |
|  | Negative class (non-ALS) | False Positive (FP) | True Negative (TN) |

Table 2 presents the input data that have two classes of output. Each column represents a predicted class, while each row indicates instances in an actual class, referred to as the *ground truth*. Based on the predictions for the test data contained in the confusion matrix, the accuracy of the classifier is evaluated. The overall accuracy 𝒜 of the classifier is defined by

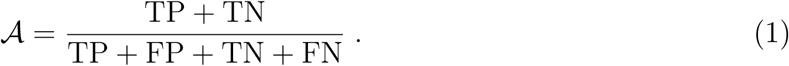

Instead of the overall accuracy, one may use another quantity called *recall* or true positive rate (TPR) ℛ^+^, which is a measure of ALS cases that are classified correctly. It is defined by

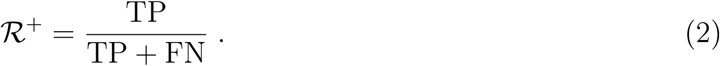

Another quantity that is used is precision 𝒫, which is indicative of the fraction of the true ALS cases among those classified as ALS, and is defined by

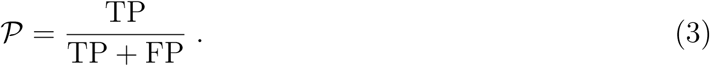

When analyzing imbalanced datasets, it is important to not only have high overall accuracy 𝒜 for the classifier, but also high TPR ℛ^+^ for the minority class. Another measure of an accurate classifier is the so-called F-measure *F*_*m*_,

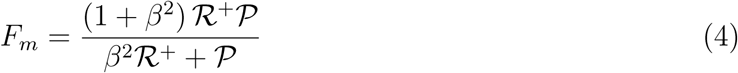

where *β* is a tunable parameter that measures relative importance of ℛ^+^ and 𝒫. Thus, *F*_*m*_ indicates that a good classifier should have both high ℛ^+^ and 𝒫. In our study we assumed that *β* = 0.2, implying that

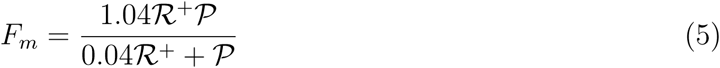

Varying *β* enables one to place more emphasis on either the TPR or the precision. The choice of *β* = 0.2 was intended to attribute more weight to the ALS data, hence ensuring that their accurate classification is more important.

## V. RESULTS

The datasets were divided between the training data (about 85% of the total) and the test data (the remaining 15%). The training data themselves were divided into training and validation set (with proportions, respectively, of 70 - 30%). The computational algorithm never “sees” or uses the test data during the training. Since we train different m odels, based o n the learning rate and several ratios of validation data, to achieve high accuracy, overfitting does not occur.

All the datasets were compared with each other based on the same 24 variables, which were SIFT_score, Polyphen2_HDIV_score, Polyphen2_HVAR score, LRT_score, MutationTaster_score, MutationAssessor_score, FATHMM_score, PROVEAN_score, VEST3_score, MetaSVM_score, MetaLR_score, CADD_raw, CADD_phred, DANN_score, fathmm.MKL_coding_score, integrated_- fitCons_score, integrated_confidence value, GERP.. _RS, phyloP7way_vertebrate, phyloP20way_- mammalian, phastCons7way_vertebrate, phastCons20way_mammalian, SiPhy_29way_logOdds, and REVEL. The training data for the algorithm contained nearly 85 percent of the datasets, with the rest used for testing the trained ML algorithm. The three most important parameters based on the confusion matrix for the training and testing the five datasets are presented in Table I.

**Table I.** Evaluation of the classifier with training and test data

| | $\mathcal{A}$ | $\mathcal{R}^+$ | $\mathcal{P}$ | F-measure |
| --- | --- | --- | --- | --- |
| Dataset A | 0.98 | 1 | 0.3 | 0.917 |
| Dataset B | 0.972 | 1 | 0.5 | 0.962 |
| Dataset C | 0.965 | 1 | 0.545 | 0.968 |
| Dataset D | 1 | 1 | 1 | 1 |
| Dataset E | 0.928 | 1 | 0.666 | 0.981 |

Since it is crucial to correctly classify the ALS data - the minority class - the true positive rate (TPR) ℛ^+^ is the most important parameter in the study, because it indicates that the trained ML algorithm better classifies the minority class, whereas 𝒜 is indicative of the overall accuracy of the classifier. In addition, the known causal mutations from the ALS database also emerged from our analyses.

To better evaluate the performance of RUSBoost, an analysis of the effect of the size of the datasets *A* − *E* on the accuracy of the trained ML algorithm for classifying majority class (non-ALS) was carried out for all the five datasets. We also resized the test dataset to be around 20 percent of the main set. Figure 1 presents the effect of the size of the non-ALS dataset on the three measures of the accuracy of the predictions listed in Table I. We first note that trends for dataset *E* is slightly different from those of the other four datasets, which is due to the size of the dataset that is smaller than the other four.

**Figure 1:**
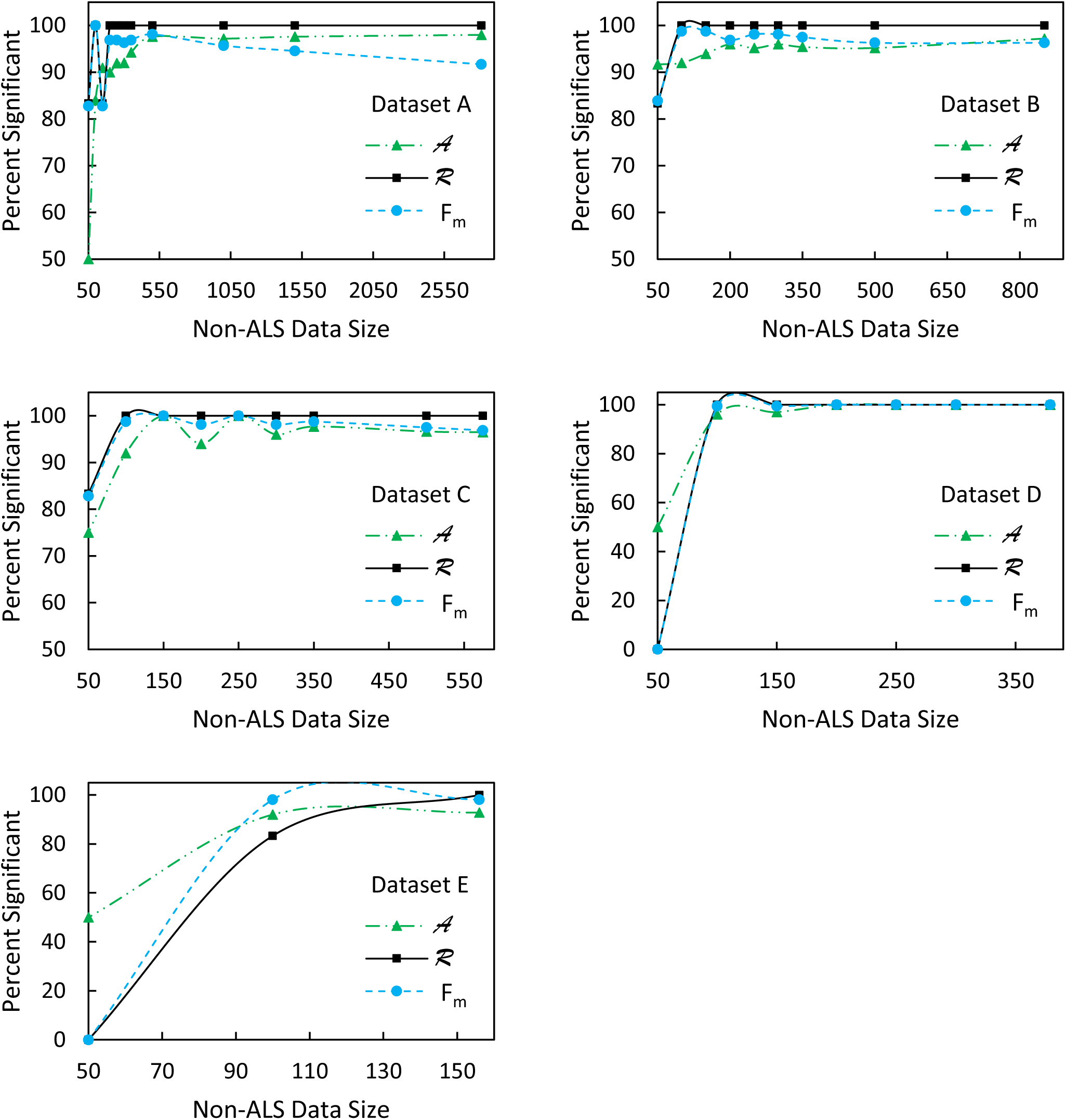
The effect of Non-ALS data size on percentage significance.

Remarkably, the accuracy ℛ^+^ of the predictions for the ALS variants is 100 percent. Without using the aforementioned dataset for a single “case” patient sample from the Coriell Institute, the overall accuracy 𝒜 of the algorithm is between 92.8 and 98 percent. We tested the network - trained with the five aforementioned datasets A, B, C, D and E - with the Coriell Institute dataset for a single patient. The ML algorithm was therefore utilized as a means of additional verification of the mutation being causal. The overall accuracy of the classification for the new dataset tested by the network was 99.18, 99.19, 99.18, 99.24, 99.31 percent for, respectively, the datasets A, B, C, D, and E.

Note that since the algorithm classifies the ALS variants with 100 percent accuracy, the overall accuracy is essentially the accuracy of the predictions for the non-ALS variants, which is also excellent. The general trends in Figure 1 suggest that, at the beginning, the rate of increase in the accuracy is very steep, but it slows down when the size of the datasets becomes relatively large. By increasing the size of the non-ALS datasets, the classifier’s overall accuracy 𝒜 and the F-measure *F*_*m*_ both increase up to a certain data size. With further increase in the dataset size, however, *F*_*m*_ and 𝒜 for the trained ML algorithm level off, and may even decrease slightly. Thus, it is crucial to select the size of the datasets carefully, in order to have the highest overall accuracy 𝒜, since it represents the accuracy of classifying the majority class, the non-ALS variants.

To identify which of the 24 variables have the strongest effect on the classifier of the ALS and non-ALS variants, we used two methods. First, we used Parallel Coordinates Plots (PCPs) that are the ideal tool for plotting multivariate data, as they allow comparing the many variables together and seeing the possible relationships between them. In a PCP each variable has its own axis, and all the axes parallel to each other. Each axis may have a distinct scale, because each variable may have a different unit of measurement. Alternatively, all the axes can be normalized to keep all the scales on all the axes uniform. Values of the variables are plotted as a series of lines that connected across all the axes, implying that each line is a collection of points placed on each axis that have all been connected together. The plot is made of the median values for each variable.

Figure 2 shows how the RUSBoost classifier has distinguished ALS from the non-ALS variants during the training process. We interpret Figure 2 as implying that the mutation assessors, FATHMM_score, PROVEAN_score, Vest3_score, CADD_phred, DANN_score, meta-SVM_score, metaLR, and REVEL are the most important variables for separating the two classes.

**Figure 2:**
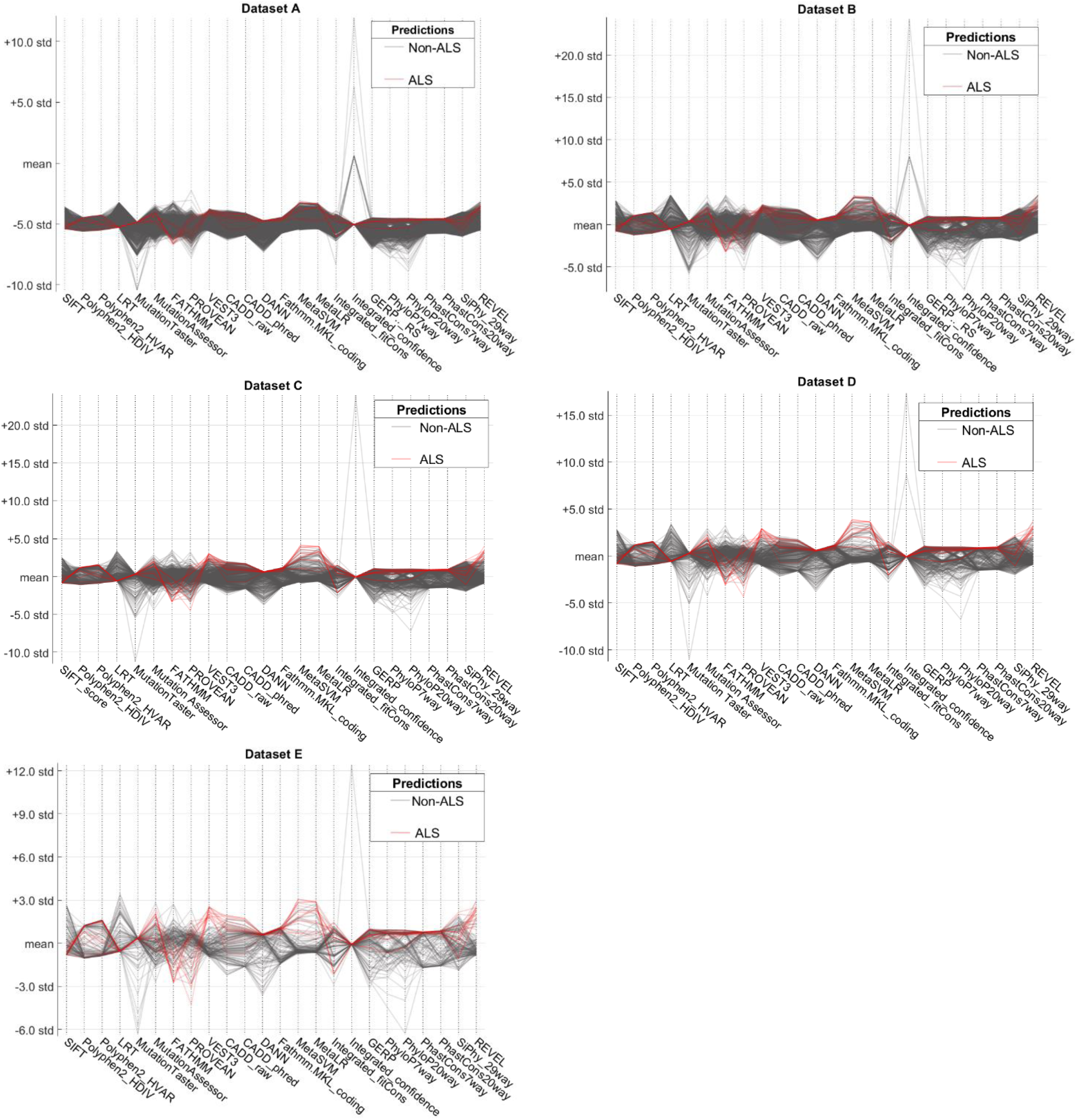
Parallel Coordinates Plot.

We also used saliency maps, usually rendered as heatmaps, for the training data, where being “hot” means being in the regions that strongly influence the algorithm’s final classification, and are particularly helpful when the algorithm incorrectly classifies a certain data point or dataset, because one can look at the input features that led to that classification. Figure 3 presents the resulting maps, showing the increase or decrease in the value of the variables for a patient. Larger values indicate that they have more influence on the final classification. Consistent with Figure 2, Figure 3 indicates that variables 7, 8, 9, 11, 12, 19, and 24, which correspond to FATHMM_score, PROVEAN_score, VEST3_score, CADD_phred, DANN_score, phyloP7way_- vertebrate, and REVEL, are the most influential mutation assessors for distinguishing the two classes of data. Thus, overall, the algorithm has identified 9 variables of mutation assessors as being the most influential.

**Figure 3:**
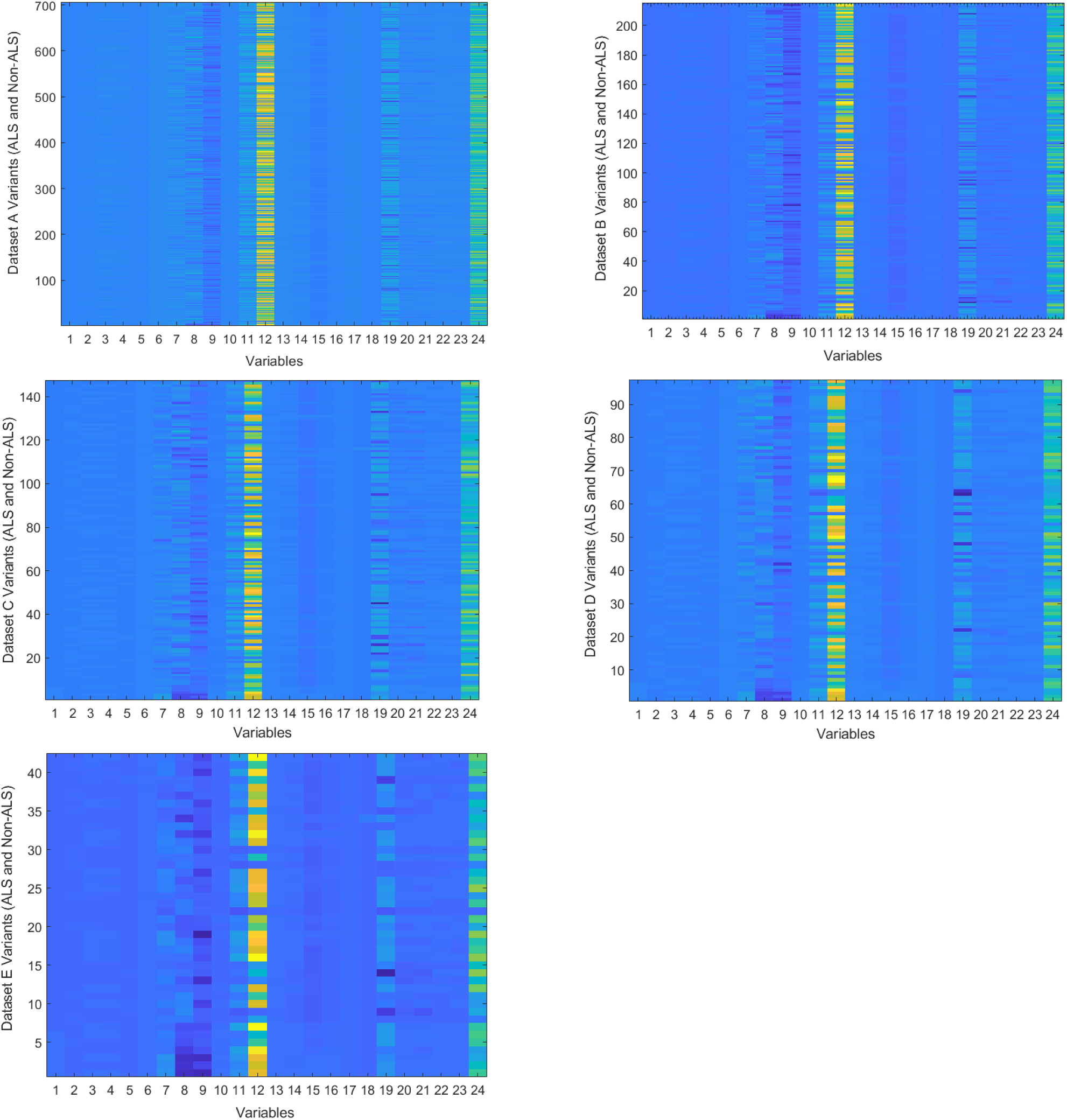
Saliency maps (heatmaps) for 24 variables in the five datasets, used for testeing the trained algorithm.

## VI. SUMMARY

A machine-learning method was proposed for classifying ALS–causative and non–ALS–causative variants based on 24 variables in 5 distinct datasets. The method predicts the ALS variants with 100 percent accuracy, while its accuracy for the non-ALS variants is up to 99.31 percent. The classifier also identifies the nine most influencial mutation assessors that help distinguishing the two classes from each other, which are FATHMM_score, PROVEAN_score, Vest3_score,

CADD_phred, DANN_score, meta-SVM_score, phyloP7way_vertebrate, metaLR, and REVEL. Therefore, they may be used in future studies to reduce the time and cost of collecting data and carrying out experimental tests, as well as in studies that are focused on the recognized assessors.

